# Heterologous expression of a lycophyte protein enhances angiosperm seedling vigor

**DOI:** 10.1101/2022.05.06.490942

**Authors:** Samuel W.H. Koh, Carlisle S. Bascom, Eduardo Berenguer, Gwyneth Ingram, Mark Estelle, Christian S. Hardtke

## Abstract

Seedling vigor is a key agronomic trait that determines juvenile plant performance. Angiosperm seeds develop inside fruits and are connected to the mother plant through vascular tissues. Their formation requires plant-specific genes, such as *BREVIS RADIX (BRX)* in *Arabidopsis thaliana* roots. BRX family proteins are found throughout the angiosperms but also occur in non-vascular bryophytes and non-seed lycophytes. They consist of four conserved domains, including the tandem “BRX-domains”. We found that bryophyte or lycophyte BRX homologs can only partially substitute for Arabidopsis BRX (AtBRX) because they miss key features in the linker between the BRX-domains. Intriguingly however, expression of a BRX homolog from the lycophyte *Selaginella moellendorffii* (SmBRX) in *A. thaliana* wildtype background confers robustly enhanced root growth vigor that persists throughout the life cycle. This effect can be traced back to a substantial increase in seed and embryo size, and can be reproduced with a modified, “SmBRX-like” variant of AtBRX. Our results thus suggest that BRX variants could serve as biotechnological tools to boost seedling vigor and shed light on the activity of ancient, non-angiosperm BRX family proteins.

## Introduction

Plant evolution is marked by major transitions that have culminated in the angiosperms, the flowering seed plants that dominate the extant terrestrial biosphere (Amborella Genome, 2013; Rensing, 2020; Spencer et al., 2021). Key inventions include the evolution of vascular tissues, which separates lycophytes from the simpler bryophytes; enclosure of the embryo in a seed, which separates spermatophytes from lycophytes; and protection of the seeds inside fruits, which separates angiosperms from gymnosperms. The development of such evolutionary novelties often entails plant-specific gene families (Armisen et al., 2008; Guo, 2013; Jiao et al., 2020; Pfannebecker et al., 2017; Rensing, 2020). Among them, the *BREVIS RADIX (BRX)* gene family comprises five members in the angiosperm model plant *Arabidopsis thaliana, AtBRX* and its homologs *AtBRX-LIKE (AtBRXL) 1-4* (Beuchat et al., 2010a; Briggs et al., 2006). The encoded BRX family proteins consist of four distinct, highly conserved domains that are not found outside the green lineage (Briggs et al., 2006; Koh et al., 2021). They include the signature tandem “BRX-domains”, which are connected by a linker of variable sequence and size (Koh et al., 2021). All BRX family proteins monitored to date are primarily plasma-membrane-associated and display polar cellular localization (Bringmann and Bergmann, 2017; Koh et al., 2021; Marhava et al., 2020; Marhava et al., 2018; Rowe et al., 2019; Scacchi et al., 2009). The biological functions of *BRX* family genes are diverse and point to sub-functionalization of individual family members (Koh et al., 2021; Li et al., 2019; Marhava et al., 2020; Rowe et al., 2019; Zhang et al., 2021). For instance, the best-characterized members, *AtBRX* and *AtBRXL2*, are interchangeable in stomata development (Rowe et al., 2019) but not in root protophloem development (Koh et al., 2021; Marhava et al., 2020), and this unequal redundancy has recently been associated with differences in protein behavior (Koh et al., 2021; Marhava et al., 2020). In the root, AtBRX guides the progression of protophloem sieve element differentiation by modulating the local trans-cellular flux of the phytohormone, auxin (Marhava et al., 2018; Moret et al., 2020). In *brx* loss-of-function mutants, protophloem differentiation is thus impaired and consequently, root growth vigor is strongly reduced (Anne and Hardtke, 2017; Moret et al., 2020; Mouchel et al., 2004; Rodrigues et al., 2009). AtBRX protein is polarly localized at the rootward end of developing protophloem sieve elements, where it interacts with an AGC-type kinase regulator of the auxin transport machinery in an intricate, feedback-regulated “molecular rheostat” (Aliaga Fandino and Hardtke, 2022; Bassukas et al., 2021; Marhava et al., 2018). A key feature of this molecular rheostat is the auxin-responsive plasma-membrane-dissociation of AtBRX (Marhava et al., 2018; Scacchi et al., 2009), which is a quantitative determinant of BRX family protein activity in the developing protophloem context (Koh et al., 2021; Marhava et al., 2020). It has been mapped to AGC kinase target phosphosites, including a key site in the linker sequence, which are present in AtBRX but absent in AtBRXL2 (Koh et al., 2021). Engineering these sites into AtBRXL2 renders the modified protein auxin-responsive and augments its biological activity in the protophloem (Koh et al., 2021). Conversely, in a BRX family protein from the lycophyte *Selaginella moellendorffii* (SmBRX), the phosphosites are missing and the linker is much shorter than in *A. thaliana* BRX family proteins (Koh et al., 2021). Thus, SmBRX plasma-membrane-association is not auxin-responsive and can only partially rescue the protophloem differentiation defects of *brx* loss-of-function mutants (Koh et al., 2021). In summary, the available data suggest that sub-functionalization of BRX family proteins is at least in part determined by the sequence of the linker between the tandem BRX-domains.

Despite their incapacity to fully complement the *brx* mutant, both AtBRXL2 and SmBRX can confer significant rescue of root growth vigor when expressed under control of the protophloem-specific *AtBRX* promoter (Beuchat et al., 2010a; Briggs et al., 2006; Koh et al., 2021). This might reflect the significant yet partial rescue of protophloem defects, which manifests for instance in a strongly reduced proportion of seedlings that display visibly impaired differentiation in both sieve element strands (Koh et al., 2021). Nevertheless, compared to other BRX family proteins that lack the key AGC kinase target phosphosite in the linker and were monitored previously (Beuchat et al., 2010a; Briggs et al., 2006; Marhava et al., 2020), the rescue of *brx* root growth vigor obtained with SmBRX was remarkable and statistically indistinguishable from the Columbia-0 (Col-0) wildtype control (Koh et al., 2021). Here, we explored this phenomenon in detail and found that SmBRX expression in *A. thaliana* wildtype substantially enhances seed vigor.

## Results

### The linker between the BRX-domains determines BRX protein family sub-functionalization

Alignment of 300 full length *bona fide* BRX family proteins retrieved from across the green lineage (One Thousand Plant Transcriptomes, 2019) shows that the key AGC target phosphosite (corresponding to S228 in AtBRX) is embedded in a 20 amino acid motif within the linker (SAXXSPVTPPLXKERLPRNF) that is conserved in the vast majority of BRX family proteins (Dataset S1 and Figure S1). Although the phosphosite serine shows the highest level of conservation (89%) within this motif, the AGC kinase consensus (R[D/E]S) is only present in a subset of ca. 10% of BRX family proteins that are all exclusively from angiosperms. Moreover, in AtBRX and its interchangeable homolog AtBRXL1 (Briggs et al., 2006; Koh et al., 2021), the R[D/E]S site is present but the motif is only partly conserved. Finally, the motif is notably absent from all lycophyte and bryophyte BRX family proteins examined.

To further determine the functional relevance of this region, we chose to investigate a few representative BRX family proteins from different phylogenetic branches that display a combination of linker features. First, we identified two *BRX* family genes in the basal angiosperm, *Amborella trichopoda* (*AmbBRXL1* and *AmbBRXL2*). Both encoded proteins have linkers of a size comparable to *A. thaliana* BRX family proteins (125 and 120 amino acids, respectively), and in both the 20 amino acid motif is conserved. However, only AmbBRXL2 carries the R[D/E]S consensus phosphosite (Figure S1). When expressed under control of the *AtBRX* promoter, an AmbBRXL1-CITRINE fusion protein codon-optimized for *A. thaliana* at best partially complemented the root growth (Figure S2A) or protophloem defects (Figure S2B) of *A. thaliana brx* mutants. By contrast, a codon-optimized AmbBRXL2-CITRINE fusion protein fully complemented all *brx* mutant defects (Figure S2C and D), and consistently, AmbBRXL2, but not AmbBRXL1, displayed auxin-induced plasma-membrane-dissociation (Figure S2K-M). These functional assays reiterate the importance of the S228 phosphosite for AtBRX-like activity.

Next, using the same complementation approach, we monitored three BRX family proteins identified in bryophytes, the two proteins identified in the *Physcomitrium patens* genome (PpBRXL1 and PpBRXL2), and the single protein found in the *Marchantia polymorpha* genome (MpBRXL1). All three proteins lack the R[D/E]S phosphosite as well as the 20 amino acid motif, but whereas PpBRXL1 and PpBRXL2 linker sizes are comparable to *A. thaliana* BRX family proteins (127 and 122 amino acids, respectively), the MpBRXL1 linker is considerably shorter (61 amino acids) (Figure S1). As expected, all three proteins only partially rescued root growth vigor or protophloem defects of *brx* mutants (Figure S2E-J), and none of them was auxin-responsive (Figure S2N-P). Since all five proteins assayed displayed protophloem-specific expression and polar localization similar to AtBRX (Figure S2Q-U), we conclude that the linker sequence between the BRX-domains is a major determinant of specific activity, consistent with previous findings (Beuchat et al., 2010a; Koh et al., 2021); and that the entire functional spectrum represented by AtBRX is only contained in a subgroup of angiosperm BRX family proteins.

### Seedling growth vigor is enhanced by heterologous SmBRX expression

Our assays of bryophyte BRX family proteins reiterate that the linker region confers the functional features that make AtBRX unique and are required for proper protophloem sieve element differentiation (Koh et al., 2021). Notably however, neither of the three bryophyte proteins consistently conferred the essentially wildtype root growth vigor observed previously in complementation experiments with SmBRX (Koh et al., 2021), which has the shortest (38 amino acids) linker identified so far and lacks any of the conserved linker sequences recognizable in most other BRX family proteins (Dataset S1). To further explore this phenomenon, we expressed SmBRX-CITRINE fusion protein under control of the *AtBRX* promoter in wildtype background. Corroborating its growth-promoting effect, these transgenic lines displayed significantly increased growth vigor that manifested in ca. 25% longer roots in 7-day-old seedlings (Figure 1A). Such root growth promotion was neither observed in similar experiments with AmbBRXL1, MpBRXL1 and AtBRXL1 (Figure 1B-D), nor upon copy number increase of AtBRXL1 or AtBRXL2 (Figure 1E and F), nor upon ectopic overexpression of AtBRX or AtBRXL2 under control of the ubiquitous *35S* promoter (Figure 1G and H). To exclude that the growth-promoting property of SmBRX was due to its codon-optimization for *A. thaliana*, we introduced a similar construct expressing a non-codon-optimized version (SmBRX^nop^) into *brx* mutants and Col-0 wildtype. Similar to codon-optimized SmBRX, this protein displayed sieve element-specific expression and polar localization (Figure S2V). Again, we observed partial complementation of protophloem defects and wildtype-level root growth vigor in *brx* (Figure S3A and B), and larger-than-wildtype root growth vigor in Col-0 (Figure S3C). Conversely, a codon-optimized version of AtBRX (AtBRX^opt^) essentially behaved like the wildtype protein (Figure S3D-F). In summary, these experiments suggest that structural features of the SmBRX protein are responsible for its growth-promoting property.

**Figure 1.**
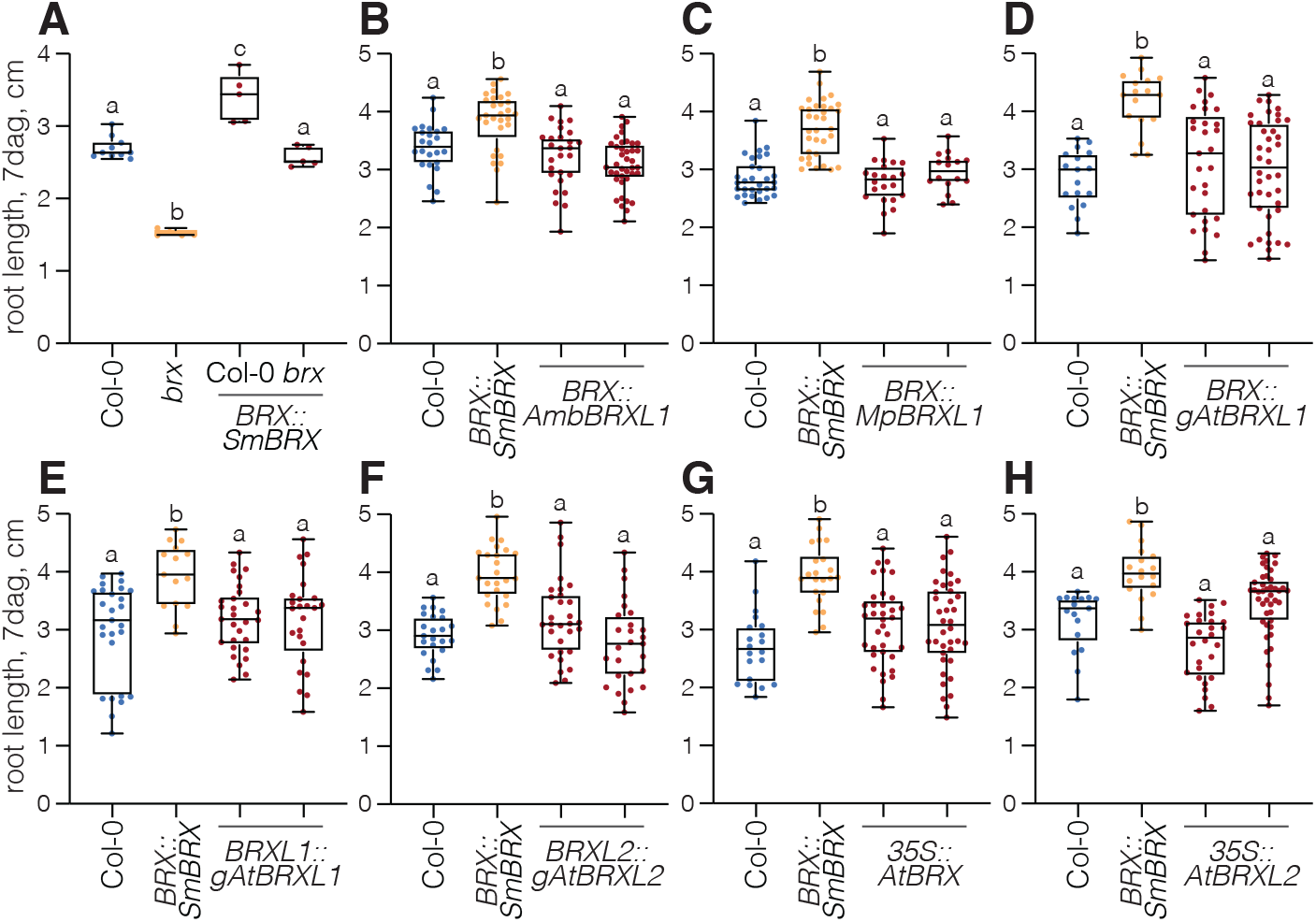
Heterologous expression of the *Selaginella moellendorffii* BRX family protein enhances root growth vigor. (A) Average root length of 7-day-old seedlings from Col-0 wildtype, *brx* mutant, and transgenic lines expressing the *S. moellendorffii BRX* homolog under control of the *A. thaliana BRX* promoter (*BRX::SmBRX*). n=5-10 independent (transgenic) lines and experiments. (B-D) Root length of 7-day-old seedlings from Col-0, *brx*, and two representative independent transgenic lines each, expressing the indicated BRX family proteins under control of the *A. thaliana BRX* promoter in Col-0. (B) n=24-39 roots; (C) n=17-33 roots; (D) n=17-40 roots. (E-F) Root length of 7-day-old seedlings from Col-0, *brx*, and two representative independent transgenic lines with a dosage increase in the indicated *A. thaliana BRX* family gene in Col-0. (E) n=15-29 roots; (F) n=23-30 roots. (G-H) Root length of 7-day-old seedlings from Col-0, *brx*, and two representative independent transgenic lines each, ectopically over-expressing the indicated *A. thaliana* BRX family protein under control of the constitutive *35S* promoter in Col-0. (G) n=20-38 roots; (H) n=17-44 roots. Box plots display 2nd and 3rd quartiles and the median, bars indicate maximum and minimum. All BRX family proteins were expressed as C-terminal CITRINE fusions. Statistically significant different samples (lower case letters) were determined by ordinary one-way ANOVA.

Next, we tested whether SmBRX mediates enhanced root growth across a range of conditions. We found that both wildtype and SmBRX transgenics responded to variations in sucrose concentration in the media, but in all conditions, the SmBRX transgenics displayed longer roots than wildtype (Figure 2A-C). In tendency, the difference in growth vigor was even amplified in the absence of sucrose (Figure 2A). Likewise, SmBRX transgenics maintained their growth advantage in various adverse conditions, such as absence of any nutrients (Figure 2D), sub-optimal pH (Figure 2E), or challenge by peptides that suppress protophloem formation (Figure 2F). Finally, although Col-0 seeds are essentially non-dormant, we stimulated germination by application of gibberellic acid to exclude that the root growth differences could result from a premature germination of SmBRX trangenics (Figure 2G). In summary, our experiments suggest that heterologous expression of SmBRX in *A. thaliana* results in a robust increase in seedling growth vigor.

**Figure 2.**
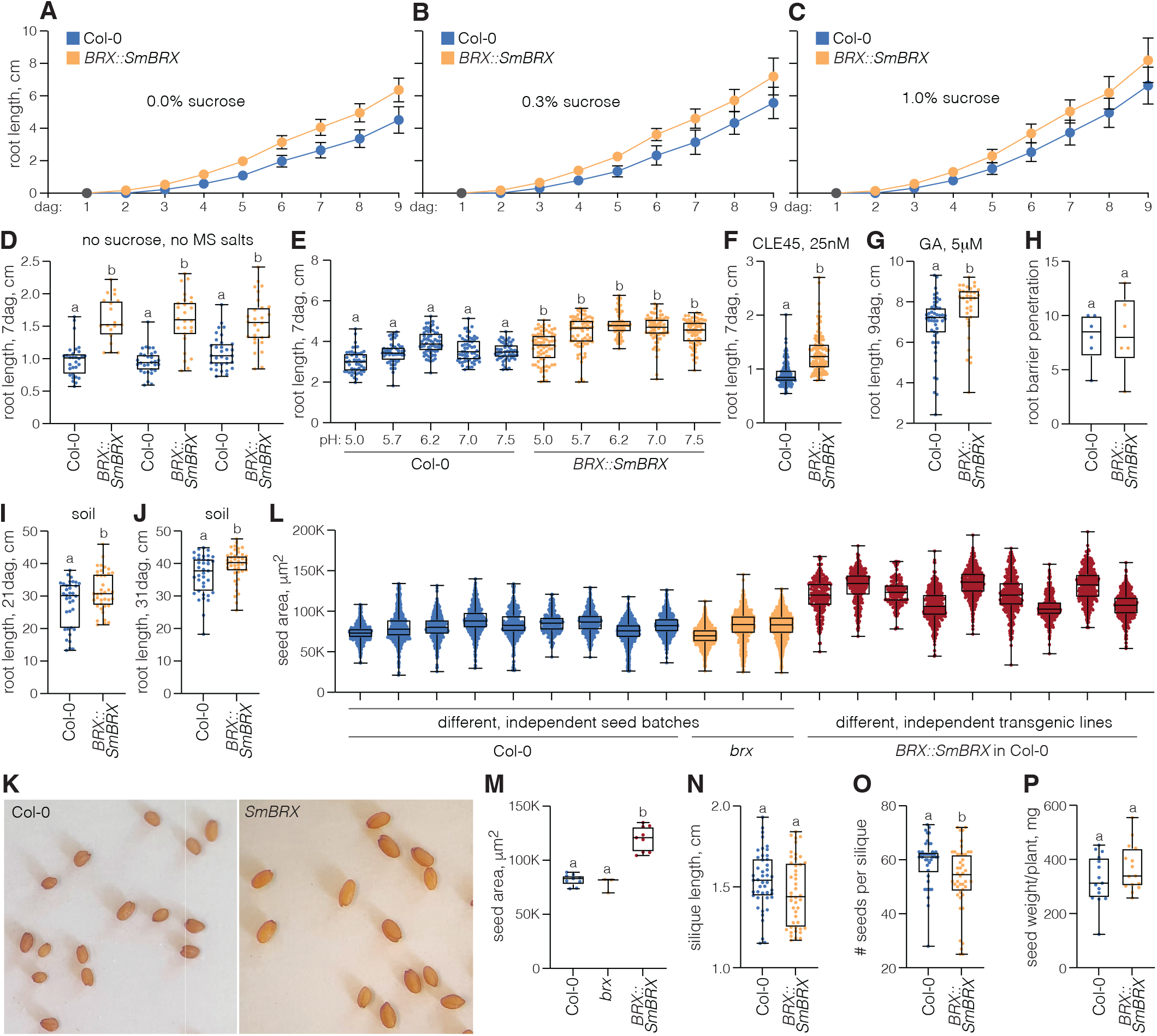
Enhanced seedling vigor in lines expressing SmBRX is robust. (A-C) Root growth progression in Col-0 wildtype and transgenic lines expressing the *S. moellendorffii BRX* homolog under control of the *A. thaliana BRX* promoter in Col-0 (*BRX::SmBRX*), on media with different sucrose supplements. n=30-51 roots per time point. Error bars indicate standard deviation. (D) Root length of 7-day-old seedlings from Col-0 and *BRX::SmBRX* plants grown on plain agar media, three independent seed batches each. n=19-32 roots. (E-G) Root length of 7- or 9-day-old seedlings from Col-0 and *BRX::SmBRX* plants, grown on media adjusted for different pH (E) or supplemented with CLE45 peptide (F) or gibberellic acid (G). (E) n=51-74 roots; (F) n=127-159 roots; (G) n=35-50 roots. (H) Number of roots that successfully penetrated a dense ca. 2 cm layer of pebbles embedded in agar media at 10 days after germination. n=6 replicates, n=20 plants per replicate. (I-J) Root length of 21- or 31-day-old Col-0 and *BRX::SmBRX* plants, grown in soil in individual rhizotrons. n=36-42 roots. Note that rhizotron length was 50 cm and *BRX::SmBRX* plants in (J) had frequently reached the bottom of the set up box. (K) Flatbed scanner images of seeds obtained from greenhouse-grown Col-0 and *BRX::SmBRX* plants, equal scale. (L) Projected area of Col-0, *brx*, and *BRX::SmBRX* seeds in flatbed scanner images, for seed batches harvested from independent lines at different times. n=357-628 seeds. (M) Projected area of Col-0, *brx*, and *BRX::SmBRX* seeds in flatbed scanner images, averages per genotype for the data shown in (L). n=3-9. (N-P) Silique length (N), seed number per silique (O), and total dry seed weight per plant (P) for greenhouse-grown Col-0 and *BRX::SmBRX* plants. (N) n=45 siliques each; (O) n=41-44 siliques; (P) n=15 plants each. Box plots display 2nd and 3rd quartiles and the median, bars indicate maximum and minimum. Statistically significant different samples (lower case letters) were determined by ordinary one-way ANOVA.

### SmBRX confers increased seed and embryo size

To determine whether the growth advantage of SmBRX transgenics in tissue culture translates into the soil environment, we first tested the capacity of their roots to vertically penetrate a ca. 2 cm thick barrier of densely packed pebbles embedded in media. In this assay, SmBRX transgenics performed as well as wildtype (Figure 2H), suggesting that they maintained their capacity to navigate a complex environment. We then monitored adult root system growth in a greenhouse setting. In replicate experiments with a setup of 96 rhizotrons, SmBRX transgenics maintained their growth advantage until at least 31 days after germination. The mean difference to wildtype remained relatively constant in comparisons between 21-day-old (Figure 2I) and 31-day-old (Figure 2J) plants however (between 3-4 cm, with average wildtype length ca. 28 and 37 cm, respectively), suggesting that SmBRX transgenics do not display a permanently higher growth rate but might carry over an early advantage. Since we had excluded a causative role of premature or accelerated germination (Figure 2G), we closely inspected the seeds. Indeed, seeds from SmBRX transgenics were visibly bigger than wildtype seeds (Figure 2K). Size approximation through image analysis of flatbed scans of dried seeds confirmed this notion and its statistical significance for a sample of independent transgenic lines and wildtype seed batches harvested at different times (Figure 2L and M). By contrast, seed batches of *brx* mutants were similar in size to wildtype (Figure 2L and M). Moreover, a high resolution image analysis of thousands of individual seeds (Figure S4A and B) confirmed these results in a comparison of wildtype with independent transgenic lines that either expressed an AtBRX-CITRINE fusion protein or an SmBRX-CITRINE fusion protein under control of the *AtBRX* promoter: wildtype and AtBRX transgenics were largely similar in the distributions of surface projection (Figure 3A), maximum seed length (Figure 3B) and maximum seed width (Figure 3C), whereas the distribution of SmBRX transgenics was skewed to higher values for all parameters tested (Figure 3A-C). Based on the average length and width, we estimated that the derived idealized ellipsoid seed volume of SmBRX transgenics is increased by about 75% compared to wildtype or AtBRX transgenics (ca. 0.190 vs ca. 0.107 cubic micrometers). In summary, our data suggest that the seeds of SmBRX transgenics are substantially bigger than in wildtype. Dried *A. thaliana* seeds largely represent the mature embryo, because the endosperm is essentially consumed as embryogenesis progresses (Doll and Ingram, 2022; Lafon-Placette and Kohler, 2014). This suggested that mature SmBRX transgenic embryos are bigger than wildtype embryos, which turned out to be the case (Figure S4C and D). Finally, the observation matches with the reported expression of *AtBRX* during embryogenesis (Bauby et al., 2007; Scacchi et al., 2009).

**Figure 3.**
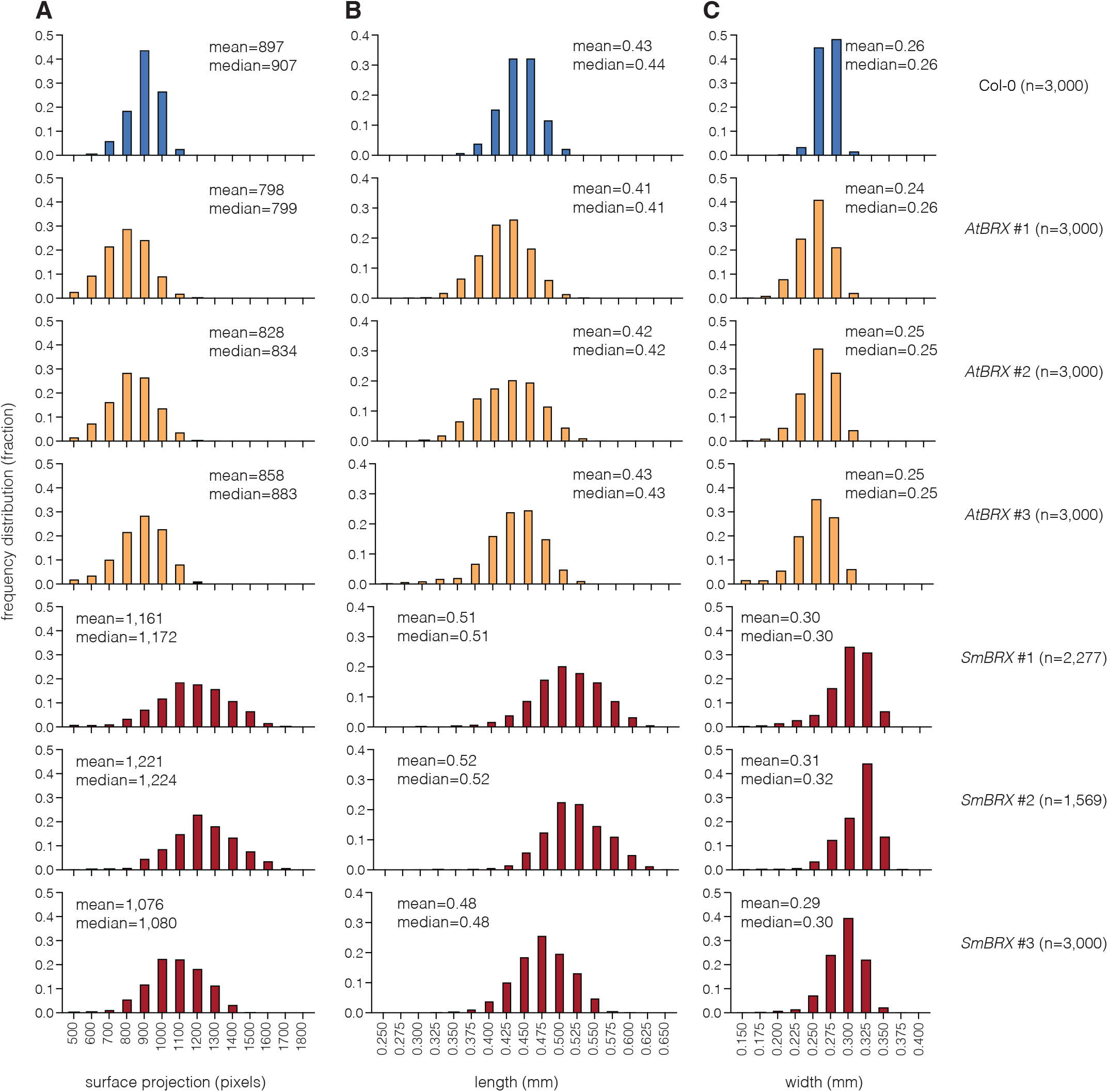
Heterologous SmBRX confers increased seed size. (A-C) High throughput, high resolution seed size parameter measurements obtained on a *Boxeed* platform for Col-0 wildtype and three independent transgenic lines each expressing either AtBRX or SmBRX under control of the *A. thaliana* BRX promoter in Col-0 background. Dry seeds were harvested from mother plants grown in parallel.

Next we asked whether the increased seed size in SmBRX transgenics is associated with tradeoffs in seed or biomass productivity? In greenhouse-grown plants, we did not observe any difference in fruit size (silique length, Figure 2N), however we observed a statistically significant, ca. 10% reduction in seed number (Figure 2O). Nevertheless, total dry seed weight per plant was similar if not in tendency higher in SmBRX transgenics (Figure 2P). Collectively, our data therefore suggest that the *SmBRX* transgene triggers a substantial increase in seed size, accompanied by a small decrease in seed number, but does not adversely affect overall plant productivity.

### An AtBRX in-frame deletion variant recapitulates SmBRX gain-of-function effects

Because SmBRX does not only diverge from AtBRX in the linker between the BRX-domains, but also in the non-conserved sequences that connect the conserved domains in the N-terminal regions, we sought to determine whether the reduced linker size of 37 amino acids was causative for the SmBRX-mediated growth promotion. To this end we engineered an AtBRX variant in which residues 219 to 266 were deleted, thereby removing the 20 amino acid motif including the RES phosphosite (Figure S1) and reducing the linker size from 93 to 45 amino acids. This “SmBRX-like” AtBRX (AtBRX^Sml^) was then introduced into Col-0 wildtype plants and *brx* mutants as a CITRINE fusion protein, expressed under control of the *AtBRX* promoter. Intriguingly, AtBRX^Sml^ transgenics displayed essentially similar features as SmBRX transgenics: in *brx* background, protophloem defects were only partially rescued (Figure 4A) although root growth vigor was similar to wildtype (Figure 4B); in wildtype background, AtBRX^Sml^ promoted root elongation as much as SmBRX (Figure 4C) and again this could be traced back to a bigger seed size (Figure 4D). In summary, these data suggest that removal of the regulatory phosphosite in conjunction with a size reduction in the linker between the BRX domains confers a dominant, growth-promoting effect on AtBRX.

**Figure 4.**
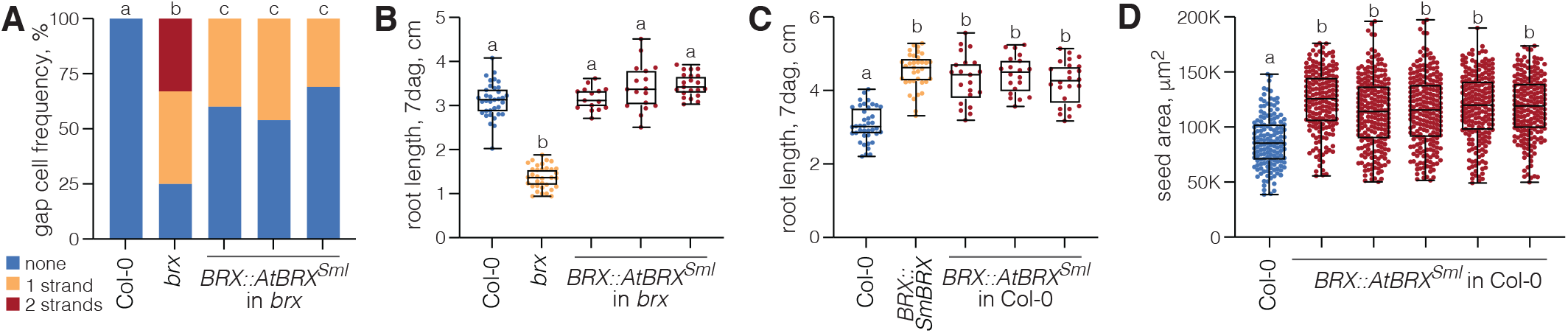
An AtBRX variant recapitulates SmBRX gain-of-function effects. (A) Quantification of protophloem sieve element differentiation defects (“gap cells”) in 7-day-old roots from Col-0 wildtype, *brx* mutant, and three independent transgenic lines expressing an AtBRX variant with an in-frame deletion of amino acids 219-266 (AtBRX^sml^) under control of the *A. thaliana BRX* promoter in the *brx* background. n=24-36 roots. (B) Root length of 7-day-old seedlings corresponding to the genotypes assayed in (A). n=15-35 roots. (C) Root length of 7-day-old Col-0 seedlings and transgenic seedlings expressing either SmBRX or AtBRX^sml^ under control of the *A. thaliana BRX* promoter in Col-0. n=20-40 roots. (D) Projected area of seeds from Col-0 and five independent transgenic AtBRX^sml^ in flatbed scanner images, for seed batches harvested from mother plants grown in parallel. n=147-216 seeds. Box plots display 2nd and 3rd quartiles and the median, bars indicate maximum and minimum. Statistically significant different samples (lower case letters) were determined by Fisher’s exact test (A) or ordinary one-way ANOVA (B-D).

## Discussion

Plants represent a variation of eukaryotic multicellularity that is distinct from animals in its unique structural, physiological and cellular characteristics. They evolved independently and it is therefore not surprising that their genomes encode proteins that are specific to the green lineage (Armisen et al., 2008; Guo, 2013; Jiao et al., 2020; One Thousand Plant Transcriptomes, 2019; Rensing, 2020). The BRX family proteins are a prime example in this context, since their highly conserved domains are not found outside plants, not even in some rudimentary form. Besides the two conserved domains in the N-terminal half of BRX proteins, the tandem BRX-domains in the C-terminal half are most prominent and can also be found in single copy in another plant-specific group of proteins that typically also contain, among others, a lipid-binding domain (Briggs et al., 2006; Furutani et al., 2020; van Leeuwen et al., 2004). The exact molecular function of the BRX-domains remains somewhat obscure, but evidence from various independent contexts suggests that they primarily mediate both homologous and heterologous protein-protein interactions and membrane attachment (Briggs et al., 2006; Furutani et al., 2020; Li et al., 2019; Marhava et al., 2020; Rowe et al., 2019; Wang et al., 2022). BRX-domains might therefore primarily represent versatile scaffolds to recruit other proteins to the plasma membrane and/or regulate their trafficking, which could explain their involvement in various processes, such as subcellular polarity establishment and maintenance or modulation of phytohormone transport (Furutani et al., 2020; Li et al., 2019; Marhava et al., 2020; Marhava et al., 2018; Moret et al., 2020; Muroyama et al., 2020; Rowe et al., 2019).

The founding member of the BRX protein family in *A. thaliana* has been identified via a loss-of-function allele, through a natural variation approach (Mouchel et al., 2004). Thus, despite the high conservation of AtBRX and its essential role in root protophloem development (Aliaga Fandino and Hardtke, 2022; Rodriguez-Villalon et al., 2014), the *AtBRX* gene is apparently dispensable in particular circumstances. Indeed, additional, extant loss-of-function alleles isolated from natural settings later on suggest that in acidic soils, where root growth is intrinsically inhibited, such alleles might even confer a competitive advantage (Gujas et al., 2012). Conversely, *AtBRX* was also identified as a quantitative trait locus that conferred slightly yet significantly increased root growth vigor (Beuchat et al., 2010a). The underlying, causative natural *AtBRX* allele encodes a hyperactive protein variant that carries a small, seven amino acid in-frame deletion in the linker between the BRX-domains (Beuchat et al., 2010a). This deletion does not affect the phosphosite that is necessary for comprehensive AtBRX function in the protophloem (Koh et al., 2021), but supports the conclusion from our current study that the length of the linker has a pivotal influence on the activity of BRX family proteins.

Among the BRX family proteins that we identified, SmBRX has the shortest linker sequence and lacks the critical phosphosite required to fully replace AtBRX in root protophloem development (Koh et al., 2021). However, additional, heterologous SmBRX expression in *A. thaliana* wildtype confers increased seed and embryo size, and subsequently enhanced seedling vigor. The engineered AtBRX^sml^ variant, which we synthesized using AtBRX as the backbone, has the same effects in both Col-0 wildtype and *brx* mutant backgrounds. This supports the idea that the unique amino acid sequence of SmBRX is not responsible for the observed gain-of-function effects, but rather structural features that depend on the distance between the two BRX-domains. Notably, this effect was observed upon expression under control of the relatively weak and (proto)phloem-specific *AtBRX* promoter (Bauby et al., 2007; Beuchat et al., 2010b; Scacchi et al., 2009). Whether the observed gain-of-function effects are directly related to the expression in the protophloem remains unclear however, because the *AtBRX* promoter confers initially wider expression during embryogenesis (Scacchi et al., 2009). Yet, this expression pattern is consistent with the increased seed and embryo size. Interestingly, the knock-out allele of *brx* in Col-0 background used in this study was originally identified because of its hypersensitive response to the phytohormone abscisic acid (Rodrigues et al., 2009). One of the various biological functions of abscisic acid is the promotion of seed dormancy, however our experiments do not support premature or accelerated germination as causative for the enhanced seedling root growth upon SmBRX or AtBRX^sml^ expression. Nevertheless, SmBRX/AtBRX^sml^-conferred gain-of-function might reflect reduced abscisic acid sensitivity, because loss-of-function mutants in the downstream abscisic acid effector ABSCISIC ACID-INSENSITIVE 5 form bigger seeds (Cheng et al., 2014; Li and Li, 2016), and because the abscisic acid antagonist, gibberellic acid, promotes post-germination root meristem growth (Achard et al., 2009; Moubayidin et al., 2010). These leads could be investigated in follow up experiments, which might also clarify why exactly SmBRX/AtBRX^sml^ seeds are bigger, since multiple interactions between seed coat, endosperm and embryo contribute to the final outcome of seed development (Doll and Ingram, 2022; Lafon-Placette and Kohler, 2014; Li and Li, 2016). Given the high conservation of BRX family proteins in both non-vascular and non-seed plants, such analyses might also aid in the characterization of their ancestral function. Irrespective of the mechanism through which SmBRX/AtBRX^sml^ gain-of-function operates, and which we did not identify here, our data suggest that BRX family protein variants can be applied as a biotechnological tool to robustly modify seedling vigor without yield penalty.

## Materials and Methods

### Sequences

The protein sequences analyzed in this paper are provided in the supplemental information, Dataset S1. The open reading frame coding sequences used for the creation of transgenes are provided in the supplemental information, Dataset S2.

### Plant material and growth conditions

The *A. thaliana* Columbia-0 (Col-0) accession was the wildtype background for all lines produced in this study. Transgenes were assayed in Col-0 or the *brx-2* mutant allele (Rodrigues et al., 2009) background. For tissue culture phenotyping assays, seeds were surface sterilized and then stratified for 2 d in the dark at ca. 4°C. Seeds were then germinated and grown in continuous white light of ca. 120μE intensity at ca. 22°C, on vertically placed Petri dishes that contained 0.5 X Murashige and Skoog (MS) media supplemented with 1% agar and 0.3% sucrose, or variations as indicated in figure panel labels. Adult plants were either monitored under controlled conditions in a walk-in chamber (16h light – 8h dark cycle with ca. 130μE light intensity, ca. 22°C, ca. 60% humidity) or a greenhouse (ca. 16h light –8 h dark cycle with variable light intensity between ca. 120-360μE, ca. 24°C, ca. 50% humidity).

### Generation of transgenic lines

Transgenic constructs for plant transformation were created in the pH7m34GW binary vector using the *Gateway*^*TM*^ cloning technology as previously described for *AtBRX* and *SmBRX* (Koh et al., 2021). Coding sequences for the BRX family proteins and variants assayed in this study are provided in the supplemental information, Dataset S2. DNA fragments for transgene construction were either obtained by gene synthesis (*AtBRX*^*opt*^, *AtBRX*^*Sml*^, *AmbBRXL1, AmbBRXL2, MpBRXL1*) (*GeneArt*^*TM*^, using the codon optimization tool where pertinent), or by RT-PCR amplification from mRNA isolated from *P. patens* or *S. moellendorffii* plants using standard procedures (*SmBRX*^*nop*^, *PpBRXL1, PpBRXL2*). All BRX family protein variants tested were expressed as C-terminal CITRINE fusion proteins to permit verification of transgene expression. All binary constructs were verified by Sanger sequencing and introduced into *Agrobacterium tumefaciens* strain GV3101pMP90 for plant transformation using the floral dip method.

### Phenotyping

For root length or seed size measurements, plates or seeds were imaged using a high resolution (1’200 dpi) flatbed scanner. Seedling root length or seed area was determined with *Fiji* image analysis software (version 2.0.1/1.53i), using suitable plug-ins. For quantification of gap cells in protophloem sieve element cell files or visualization of fluorescent protein localization, roots were imaged by confocal microscopy as described (Koh et al., 2021). High throughput and high resolution seed size measurements were performed on the *Boxeed* platform run by *Labdeers Ltd*. For trait quantification in adult *A. thaliana*, plants were grown in soil individually, in 50 cm long open-ended polyvinyl chloride tubes of 3.5cm diameter that were arranged in a 96 (8 × 12) set up. The tubes were lined with thin plastic sheets to facilitate removal of the root system and subsequent careful soil washout. Root systems were then imaged together with a ruler under a fixed distance camera set up and measured. Shoot productivity (seed number, silique length, seed weight) was monitored at end-of-life. All phenotyping assays were performed using homozygous T3 or T4 transgenic lines with verified transgene expression.

### Sequence analyses

BRX family protein sequences were retrieved from the “One KP” transcriptome dataset (One Thousand Plant Transcriptomes, 2019) and aligned and analyzed using *SnapGene* software (version 6.0.2).

### Quantification and statistical analysis

Analyses to determine statistical significance were performed in Graphpad Prism software, version 9.3.1. Specific statistical tests used are indicated in the figure legends and were always two-tailed.

## Acknowledgments

The authors would like to thank Prof. T. Beeckman for *Selaginella moellendorffii* plants, and A. Amiguet-Vercher and N. Diaz-Ardila for technical support. Author contributions: Conceptualization, S.W.H.K. and C.S.H.; Methodology, S.W.H.K. and C.S.B.; Investigation, S.W.H.K. and C.S.B.; Validation, S.W.H.K., C.S.B. and E.B.; Visualization, S.W.H.K. and C.S.H.; Writing – Original Draft, S.W.H.K. and C.S.H.; Writing – Review & Editing, S.W.H.K., C.S.B., M.E. and C.S.H.; Funding Acquisition, M.E. and C.S.H.; Resources, C.S.B.; Supervision, G.I., M.E. and C.S.H.

## Competing interests

The authors declare no competing interests.

## Funding

This study was funded by *Swiss National Science Foundation* grant 310030B_185379 awarded to C.S.H.

## Supplemental items

**Supplemental Dataset 1** Protein sequence alignment of 302 BRX family proteins retrieved from a cross the green lineage with scaffold IDs (One Thousand Plant Transcriptomes, 2019) and species names, html file.

**Supplemental Dataset 2** Open reading frame coding sequences of BRX family genes and variants used for the creation of transgenes, pdf file.

---

**Figure S1.**
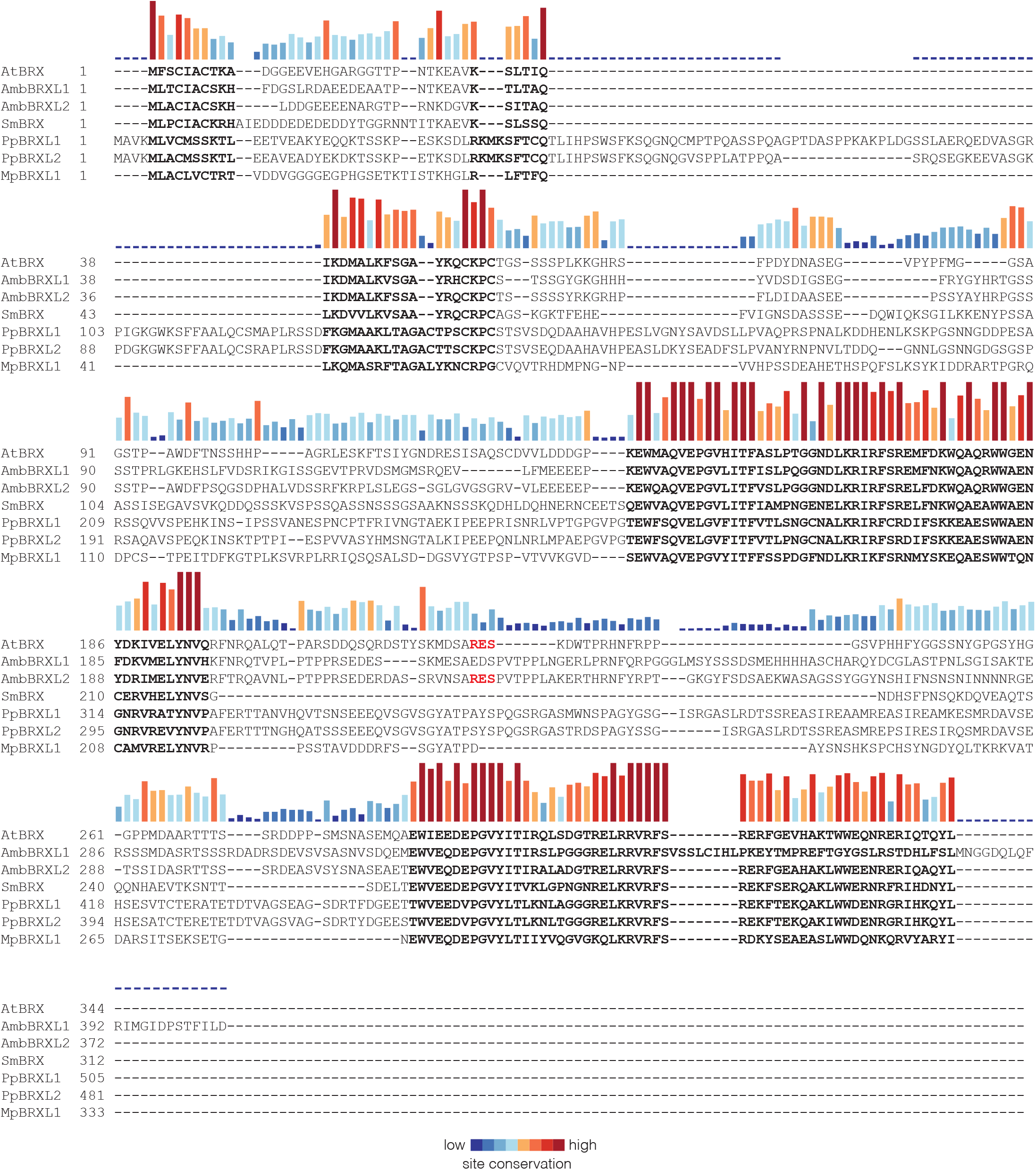
Sequence alignment. Alignment of BRX family proteins from *Arabidopsis thaliana* (AtBRX), *Amborella trichopoda* (AmbBRXLl and AmbBRXL2), *Selaginella moellendorffii* (SmBRX), *Physcomitrium patens* genome (PpBRXLl and PpBRXL2), and *Marchantia polymorpha* genome (MpBRXLl). The AGC kinase target phosphosite of AtBRX and AmbBRXL2 in the linker between the BRX-domains is highlighted in red.

**Figure S2.**
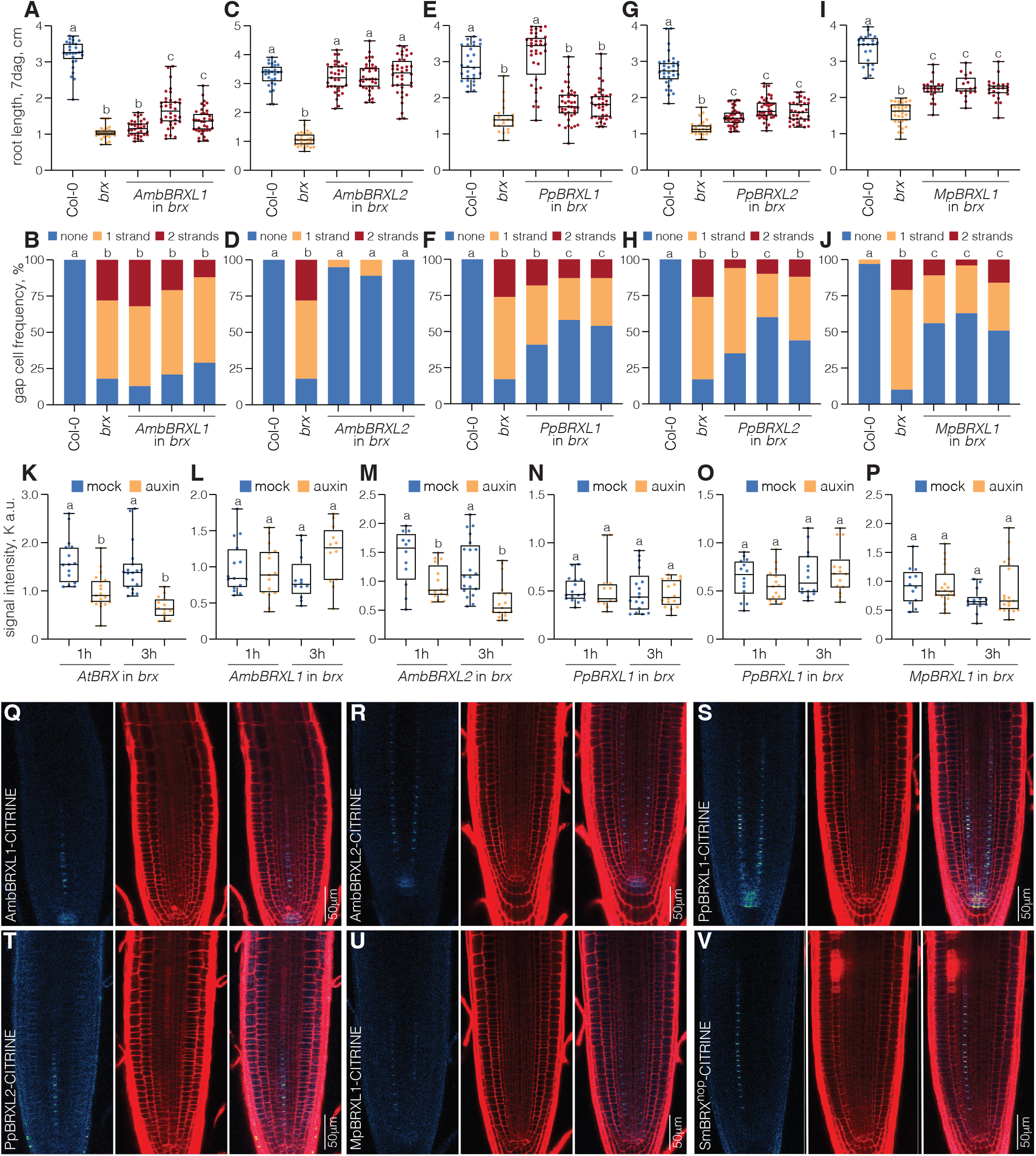
Functional assays of BRX family proteins across the green lineage. (A-J) Root length of 7-day-old seedlings and corresponding quantification of protophloem sieve element differentiation defects(“gap cells”) in 7-day-old rootsfrom Col-0 wildtype, *brx* mutant, and three representative independent transgenic lines each expressing the indicated BRX family proteins under control of the *A. thaliana BRX* promoter in *brx* background. (A) n=26-38 roots; (B) n=l 7-31 roots; (C) n=28-38 roots; (D) n=26-41 roots; (E) n=21-40 roots; (F) n=22-31 roots; (G) n=25-40 roots; (H) n=17-25 roots; (I) n=17-33 roots; (J) n=29-45 roots. (K-P) Signal intensity quantification of indicated CITRINE fusion proteins (expressed under control of the *A. thaliana BRX* promoter) at the rootward plasma membrane of developing protophloem sieve elements, lh or 3h after mock (DMSO) or auxin (10µM 1-naphthalene-acetic acid) treatment. (Q-V) Confocal microscopy images of indicated CITRINEfusion proteins (expressed under control of the *A. thafiana BRX* promoter), showing their protophloem-specific expression and polar localization (green fluorescence, left panels). Central panels show propidium iodide-stained cell wall outline (red fluorescence), right panels show the overlay. Box plots display 2nd and 3rd quartiles and the median, bars indicate maximum and minimum. Statistically significant different samples (lower case letters) were determined by Fisher’s exact test (B, D, F, H, J) or ordinary one-way ANOVA (A, C, E, G, I, K-P).

**Figure S3.**
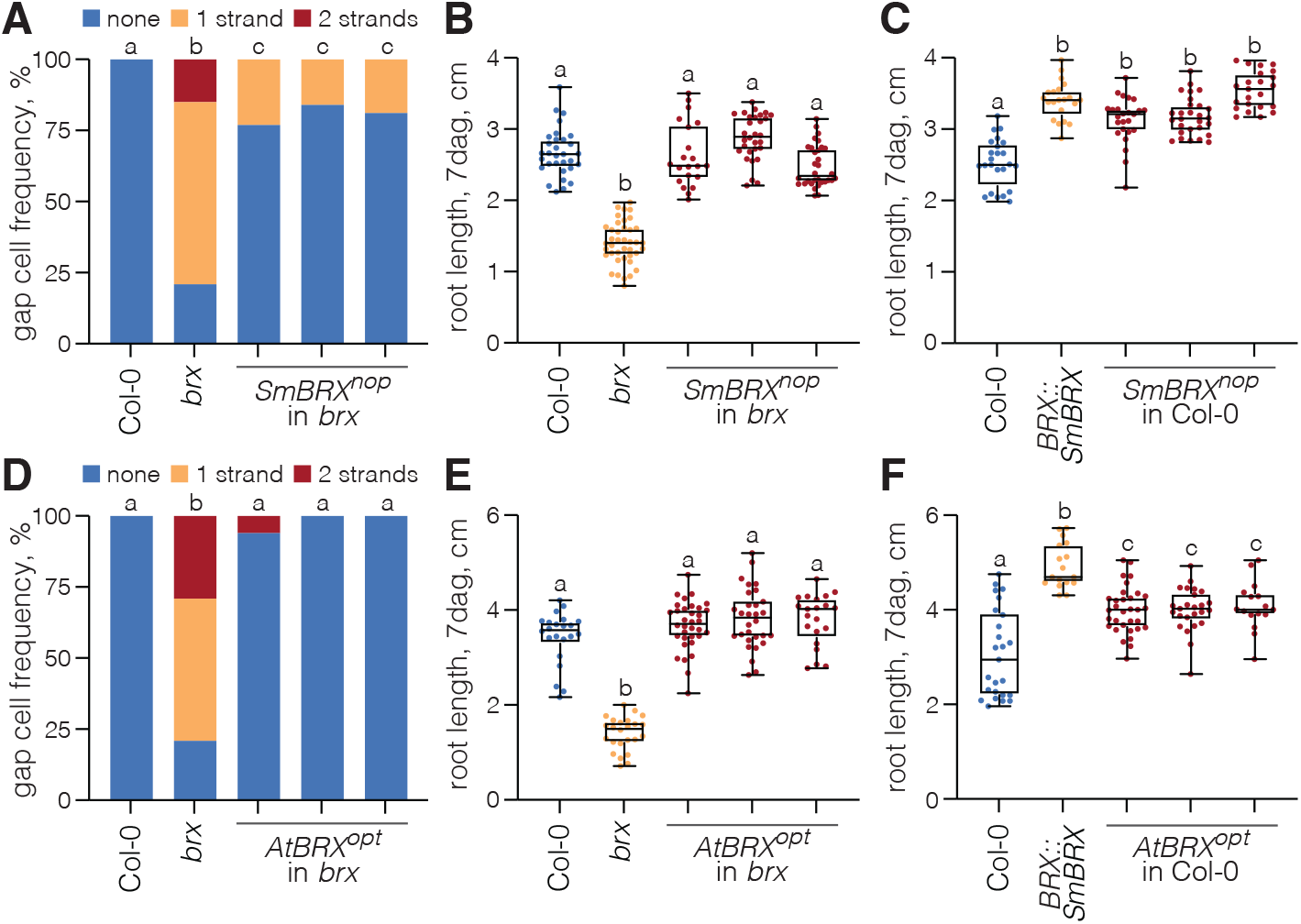
Functional assays of codon-(non)-optimized SmBRX and AtBRX variants. (A) Quantification of protophloem sieve element differentiation defects (“gap cells”) in 5-day-old roots from Col-0 wildtype, *brx* mutant, and three independent transgenic lines expressing a non-codon-optimized SmBRX variant (SmBRX^nop^) under control of the *A. thaliana BRX* promoter in the *brx* background. n = 27-48 roots. (B) Root length of 7-day-old seedlings corresponding to the genotypes assayed in (A). n = 21-40 roots. (C) Root length of 7-day-old seedlings from Col-0, *brx*, and three independent transgenic lines expressing SmBRX^nop^ under control of the *A. thaliana BRX* promoter in Col-0 background. n = 21-28 roots. (D-F) Similar to (A-C), for an *A. thaliana* codon-optimized AtBRX variant (AtBRX^opt^). (D) n = 14-27 roots; (E) n = 21-34 roots; (F) n = 16-32 roots. Box plots display 2nd and 3rd quartiles and the median, bars indicate maximum and minimum. Statistically significant different samples (lower case letters) were determined by Fisher’s exact test (A, D) or ordinary one-way ANOVA (B, C, E, F).

**Figure S4.**
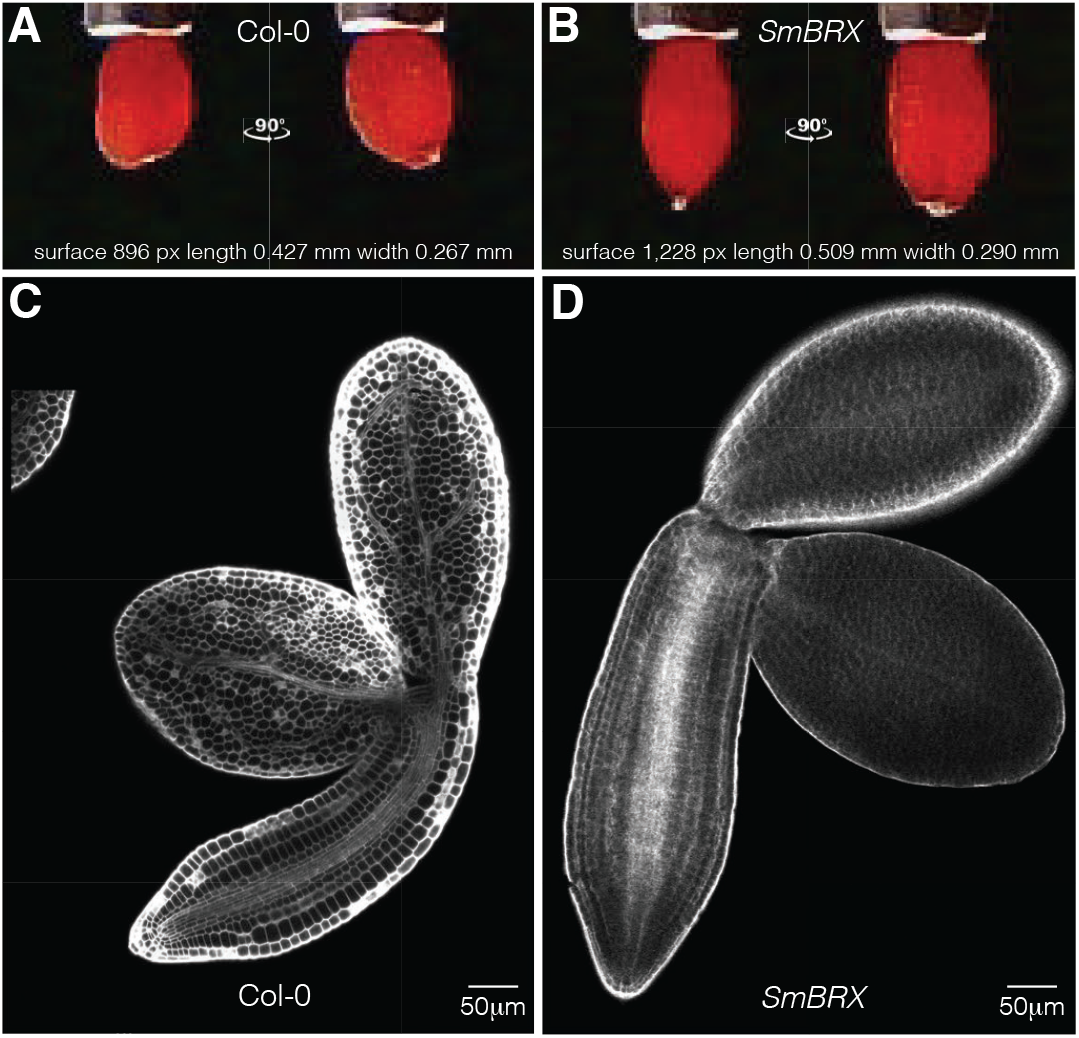
Illustration of increased seed and embryo size upon heterologous SmBRX expression. (A-B) Illustration of high resolution seed imaging with the *Boxeed* platform, for a Col-0 wildtype (A) and an SmBRX transgenic seed (B). Measured parameter values for the seeds shown are indicated. (C-D) Confocal microscopy images of calcofluor white-stained mature embryos, dissected from a Col-0 (C) or SmBRX transgenic seed (D).

## Notes

### Competing Interest Statement

The authors have declared no competing interest.

## References

Achard, P., Gusti, A., Cheminant, S., Alioua, M., Dhondt, S., Coppens, F., Beemster, G. T. and Genschik, P. (2009). Gibberellin signaling controls cell proliferation rate in Arabidopsis. Current biology : CB 19, 1188–1193.

Aliaga Fandino, A. C. and Hardtke, C. S. (2022). Auxin transport in developing protophloem: A case study in canalization. J Plant Physiol 269, 153594.

Amborella Genome, P. (2013). The Amborella genome and the evolution of flowering plants. Science 342, 1241089.

Anne, P. and Hardtke, C. S. (2017). Phloem function and development-biophysics meets genetics. Curr Opin Plant Biol 43, 22–28.

Armisen, D., Lecharny, A. and Aubourg, S. (2008). Unique genes in plants: specificities and conserved features throughout evolution. BMC Evol Biol 8, 280.

Bassukas, A. E. L., Xiao, Y. and Schwechheimer, C. (2021). Phosphorylation control of PIN auxin transporters. Curr Opin Plant Biol 65, 102146.

Bauby, H., Divol, F., Truernit, E., Grandjean, O. and Palauqui, J. C. (2007). Protophloem differentiation in early Arabidopsis thaliana development. Plant Cell Physiol 48, 97–109.

Beuchat, J., Li, S., Ragni, L., Shindo, C., Kohn, M. H. and Hardtke, C. S. (2010a). A hyperactive quantitative trait locus allele of Arabidopsis BRX contributes to natural variation in root growth vigor. Proc Natl Acad Sci U S A 107, 8475–8480.

Beuchat, J., Scacchi, E., Tarkowska, D., Ragni, L., Strnad, M. and Hardtke, C. S. (2010b). BRX promotes Arabidopsis shoot growth. New Phytol 188, 23–29.

Briggs, G. C., Mouchel, C. F. and Hardtke, C. S. (2006). Characterization of the plant-specific BREVIS RADIX gene family reveals limited genetic redundancy despite high sequence conservation. Plant Physiol 140, 1306–1316.

Bringmann, M. and Bergmann, D. C. (2017). Tissue-wide Mechanical Forces Influence the Polarity of Stomatal Stem Cells in Arabidopsis. Current biology : CB 27, 877–883.

Cheng, Z. J., Zhao, X. Y., Shao, X. X., Wang, F., Zhou, C., Liu, Y. G., Zhang, Y. and Zhang, X. S. (2014). Abscisic acid regulates early seed development in Arabidopsis by ABI5-mediated transcription of SHORT HYPOCOTYL UNDER BLUE1. Plant Cell 26, 1053–1068.

Doll, N. M. and Ingram, G. C. (2022). Embryo-Endosperm Interactions. Annu Rev Plant Biol.

Furutani, M., Hirano, Y., Nishimura, T., Nakamura, M., Taniguchi, M., Suzuki, K., Oshida, R., Kondo, C., Sun, S., Kato, K., et al. (2020). Polar recruitment of RLD by LAZY1-like protein during gravity signaling in root branch angle control. Nat Commun 11, 76.

Gujas, B., Alonso-Blanco, C. and Hardtke, C. S. (2012). Natural Arabidopsis brx Loss-of-Function Alleles Confer Root Adaptation to Acidic Soil. Current biology : CB 22, 1962–1968.

Guo, Y. L. (2013). Gene family evolution in green plants with emphasis on the origination and evolution of Arabidopsis thaliana genes. Plant J 73, 941–951.

Jiao, C., Sorensen, I., Sun, X., Sun, H., Behar, H., Alseekh, S., Philippe, G., Palacio Lopez, K., Sun, L., Reed, R., et al. (2020). The Penium margaritaceum Genome: Hallmarks of the Origins of Land Plants. Cell 181, 1097–1111 e1012.

Koh, S. W. H., Marhava, P., Rana, S., Graf, A., Moret, B., Bassukas, A. E. L., Zourelidou, M., Kolb, M., Hammes, U. Z., Schwechheimer, C., et al. (2021). Mapping and engineering of auxin-induced plasma membrane dissociation in BRX family proteins. Plant Cell 33, 1945–1960.

Lafon-Placette, C. and Kohler, C. (2014). Embryo and endosperm, partners in seed development. Curr Opin Plant Biol 17, 64–69.

Li, N. and Li, Y. (2016). Signaling pathways of seed size control in plants. Curr Opin Plant Biol 33, 23–32.

Li, Z., Liang, Y., Yuan, Y., Wang, L., Meng, X., Xiong, G., Zhou, J., Cai, Y., Han, N., Hua, L., et al. (2019). OsBRXL4 Regulates Shoot Gravitropism and Rice Tiller Angle through Affecting LAZY1 Nuclear Localization. Mol Plant 12, 1143–1156.

Marhava, P., Aliaga Fandino, A. C., Koh, S. W. H., Jelinkova, A., Kolb, M., Janacek, D. P., Breda, A. S., Cattaneo, P., Hammes, U. Z., Petrasek, J., et al. (2020). Plasma Membrane Domain Patterning and Self-Reinforcing Polarity in Arabidopsis. Dev Cell 52, 223–235 e225.

Marhava, P., Bassukas, A. E. L., Zourelidou, M., Kolb, M., Moret, B., Fastner, A., Schulze, W. X., Cattaneo, P., Hammes, U. Z., Schwechheimer, C., et al. (2018). A molecular rheostat adjusts auxin flux to promote root protophloem differentiation. Nature 558, 297–300.

Moret, B., Marhava, P., Aliaga Fandino, A. C., Hardtke, C. S. and Ten Tusscher, K. H. W. (2020). Local auxin competition explains fragmented differentiation patterns. Nat Commun 11, 2965.

Moubayidin, L., Perilli, S., Dello Ioio, R., Di Mambro, R., Costantino, P. and Sabatini, S. (2010). The rate of cell differentiation controls the Arabidopsis root meristem growth phase. Current biology : CB 20, 1138–1143.

Mouchel, C. F., Briggs, G. C. and Hardtke, C. S. (2004). Natural genetic variation in Arabidopsis identifies BREVIS RADIX, a novel regulator of cell proliferation and elongation in the root. Genes Dev 18, 700–714.

Muroyama, A., Gong, Y. and Bergmann, D. C. (2020). Opposing, Polarity-Driven Nuclear Migrations Underpin Asymmetric Divisions to Pattern Arabidopsis Stomata. Current biology : CB 30, 4467–4475 e4464.

One Thousand Plant Transcriptomes, I. (2019). One thousand plant transcriptomes and the phylogenomics of green plants. Nature 574, 679–685.

Pfannebecker, K. C., Lange, M., Rupp, O. and Becker, A. (2017). Seed Plant-Specific Gene Lineages Involved in Carpel Development. Mol Biol Evol 34, 925–942.

Rensing, S. A. (2020). How Plants Conquered Land. Cell 181, 964–966.

Rodrigues, A., Santiago, J., Rubio, S., Saez, A., Osmont, K. S., Gadea, J., Hardtke, C. S. and Rodriguez, P. L. (2009). The short-rooted phenotype of the brevis radix mutant partly reflects root abscisic acid hypersensitivity. Plant Physiol 149, 1917–1928.

Rodriguez-Villalon, A., Gujas, B., Kang, Y. H., Breda, A. S., Cattaneo, P., Depuydt, S. and Hardtke, C. S. (2014). Molecular genetic framework for protophloem formation. Proc Natl Acad Sci U S A 111, 11551–11556.

Rowe, M. H., Dong, J., Weimer, A. K. and Bergmann, D. C. (2019). A Plant-Specific Polarity Module Establishes Cell Fate Asymmetry in the Arabidopsis Stomatal Lineage. bioRxiv.

Scacchi, E., Osmont, K. S., Beuchat, J., Salinas, P., Navarrete-Gomez, M., Trigueros, M., Ferrandiz, C. and Hardtke, C. S. (2009). Dynamic, auxin-responsive plasma membrane-to-nucleus movement of Arabidopsis BRX. Development 136, 2059–2067.

Spencer, V., Nemec Venza, Z. and Harrison, C. J. (2021). What can lycophytes teach us about plant evolution and development? Modern perspectives on an ancient lineage. Evol Dev 23, 174–196.

van Leeuwen, W., Okresz, L., Bogre, L. and Munnik, T. (2004). Learning the lipid language of plant signalling. Trends Plant Sci 9, 378–384.

Wang, L., Li, D., Yang, K., Guo, X., Bian, C., Nishimura, T., Le, J., Morita, M. T., Bergmann, D. C. and Dong, J. (2022). Connected function of PRAF/RLD and GNOM in membrane trafficking controls intrinsic cell polarity in plants. Nat Commun 13, 7.

Zhang, Y., Liang, J., Cai, X., Chen, H., Wu, J., Lin, R., Cheng, F. and Wang, X. (2021). Divergence of three BRX homoeologs in Brassica rapa and its effect on leaf morphology. Hortic Res 8, 68.

